# ERM-1 phosphorylation and NRFL-1 redundantly control lumen formation in the *C. elegans* intestine

**DOI:** 10.1101/2021.09.02.458700

**Authors:** Jorian J. Sepers, João J. Ramalho, Jason R. Kroll, Ruben Schmidt, Mike Boxem

**Affiliations:** Division of Developmental Biology, Institute of Biodynamics and Biocomplexity, Department of Biology, Faculty of Science, Utrecht University, Padualaan 8, 3584 CH, Utrecht, The Netherlands; Laboratory of Biochemistry, Wageningen University & Research, Stippeneng 4, 6708 WE, Wageningen, The Netherlands

**Keywords:** ezrin, radixin, moesin, ERM-1, NHERF, EBP50, E3KARP, NRFL-1, lumenogenesis

## Abstract

Reorganization of the plasma membrane and underlying actin cytoskeleton into specialized domains is essential for the functioning of most polarized cells in animals. Proteins of the ezrin-radixin-moesin (ERM) and Na^+^/H^+^ exchanger 3 regulating factor (NHERF) family are conserved regulators of cortical specialization. ERM proteins function as membrane-actin linkers and as molecular scaffolds that organize the distribution of proteins at the membrane. NHERF proteins are PDZ-domain containing adapters that can bind to ERM proteins and extend their scaffolding capability. Here, we investigate how ERM and NHERF proteins function in regulating intestinal lumen formation in the nematode *Caenorhabditis elegans. C. elegans* has single ERM and NHERF family proteins, termed ERM-1 and NRFL-1, and ERM-1 was previously shown to be critical for intestinal lumen formation. Using CRISPR/Cas9-generated *nrfl-1* alleles we demonstrate that NRFL-1 localizes at the intestinal microvilli, and that this localization is depended on an interaction with ERM-1. However, *nrfl-1* loss of function mutants are viable and do not show defect in intestinal development. Interestingly, combining *nrfl-1* loss with *erm-1* mutants that either block or mimic phosphorylation of a regulatory C-terminal threonine causes severe defects in intestinal lumen formation. These defects are not observed in the phosphorylation mutants alone, and resemble the effects of strong *erm-1* loss of function. The loss of NRFL-1 did not affect the localization or activity of ERM-1. Together, these data indicate that ERM-1 and NRFL-1 function together in intestinal lumen formation in *C. elegans*. We postulate that the functioning of ERM-1 in this tissue involves actin-binding activities that are regulated by the C-terminal threonine residue and the organization of apical domain composition through NRFL-1.

## Introduction

The establishment of molecularly and functionally distinct apical, basal, and lateral domains is a key feature of polarized epithelial cells. The outside-facing apical domain has a different lipid and protein composition than the basal and lateral domains and is often decorated by microvilli. The specialization of the apical domain and microvilli formation requires the activities of the ezrin/radixin/moesin (ERM) family of proteins. ERM proteins consist of an N-terminal band Four-point-one/ezrin/radixin/moesin (FERM) domain that mediates binding to the plasma membrane and membrane-associated proteins, a C-terminal tail that mediates actin binding, and a central α-helical linker region (Fehon et al., 2010; McClatchey, 2014). In the cytoplasm, ERM proteins are kept in an inactive, closed, conformation that masks most of regulatory and protein interaction motifs due to an intramolecular interaction between the N- and C-terminal domains (Gary and Bretscher, 1995; Li et al., 2007; Magendantz et al., 1995; Pearson et al., 2000). Binding to the plasma membrane lipid phosphatidylinositol-(4,5) bisphosphate (PIP_2_) as well as phosphorylation of a conserved C-terminal threonine residue (T567 in ezrin) promote the transition to an open and active conformation that can link the plasma membrane to the underlying actin cytoskeleton and control the spatial distribution of protein complexes at the membrane (Simons et al., 1998; Nakamura et al., 1999; Barret et al., 2000; Fievet et al., 2004; Coscoy et al., 2002; Yonemura et al., 2002; Hao et al., 2009; Roch et al., 2010).

The ability of ERM proteins to associate with other proteins can be extended by binding to the scaffolding proteins NHERF1 and NHERF2 (Na^+^/H^+^ exchanger regulatory factors 1 and 2). NHERF1/2 were identified as co-regulators of the Na^+^/H^+^ exchanger NHE3 in kidney epithelial cells (Lamprecht et al., 1998; Weinman et al., 1993; Yun et al., 1997). Independently, NHERF1 was identified as the ERM-binding phosphoprotein 50 (EBP50), based on its ability to interact with activated ezrin and moesin (Reczek et al., 1997). NHERF1/2 are closely related proteins that contain two postsynaptic density 95/disks large/zona occludens-1 (PDZ) domains and an ERM-binding (EB) C-terminal tail that can bind to the FERM domain of active ERM proteins. Since their discovery, a large variety of NHERF1/2 interactors have been identified, including transporters like the cystic fibrosis transmembrane conductance regulator (CFTR) (Seidler et al., 2009), growth factor receptors including EGFR and PDGFR (Lazar et al., 2004; Maudsley et al., 2000), and other scaffold proteins such as the NHERF family member PDZK1 (PDZ domain containing 1) (LaLonde and Bretscher, 2009).

The functional significance of the interaction of NHERF1/2 with ERM proteins is best understood for NHERF1. In JEG3 cells, NHERF1 promotes microvilli formation or stability by acting as a linker between ezrin and PDZK1, and mice lacking either ezrin or EBP50 show similar defects in microvilli formation and organization in the intestine (Garbett et al., 2010; LaLonde et al., 2010; Morales et al., 2004; Saotome et al., 2004). In a model of MDCK cells developing into 3D cysts, a complex of NHERF1, ezrin, and Podocalyxin promotes apical identity and is required for lumen formation (Bryant et al., 2014). In a different 3D cyst model grown from Caco-2 colorectal cells, NHERF1 is similarly required for apical–basal polarization and lumen formation, but in conjunction with moesin rather than ezrin (Georgescu et al., 2014).

In addition to extending the scaffolding capacity of ERM proteins, NHERF proteins have also been reported to regulate the activity of ERM proteins. In NHERF1 knockout mice, levels of ERM proteins in membrane fractions of kidney and intestinal epithelial cells are decreased, suggesting that NHERF1 stabilizes ERM proteins at the plasma membrane (Morales et al., 2004). In *Drosophila* follicle cells, the single NHERF1/2 ortholog Sip1 is thought to promote phosphorylation and activation of Moesin through recruitment of the Ste20-family kinase Slik (Hughes et al., 2010). In an ovarian cancer cell line, depletion of NHERF1 led to reduced levels of phosphorylated ERM (pERM) upon stimulation with lysophosphatidic acid (LPA) (Oh et al., 2017). Similarly, NHERF2 was found to promote the phosphorylation of ERM in bovine pulmonary artery endothelial cells, possibly through an interaction with Rho kinase 2 (ROCK2) (Boratkó and Csortos, 2013). Finally, NHERF1 may also indirectly affect the localization of ERM proteins, by promoting the local accumulation of PIP_2_ through recruitment of lipid phosphatases or kinases (Georgescu et al., 2014; Ikenouchi et al., 2013). Thus, NHERF proteins may function both as ERM effectors and regulators.

Here, we make use of the nematode *Caenorhabditis elegans* to better understand how NHERF and ERM proteins function together to promote apical domain identity. The *C. elegans* genome encodes single orthologs of each protein family, termed NRFL-1 and ERM-1, that are highly similar in sequence and domain composition to their counterparts in other organisms (Fig. 1A; Fig. S1A). ERM-1 localizes to the apical surface of several epithelial tissues and is essential for apical membrane morphogenesis in the intestine (Göbel et al., 2004; van Fürden et al., 2004). Loss of *erm-1* in the intestine causes constrictions, loss of microvilli, severe reduction in the levels of apical actin, and defects in the accumulation of junctional proteins (Bernadskaya et al., 2011; Göbel et al., 2004; van Fürden et al., 2004). Recently, we demonstrated that the functioning of ERM-1 critically depends on its ability to bind membrane phospholipids, while phosphorylation of a C-terminal regulatory threonine residue modulates ERM-1 apical localization and dynamics (Ramalho et al., 2020).

**Figure 1.**
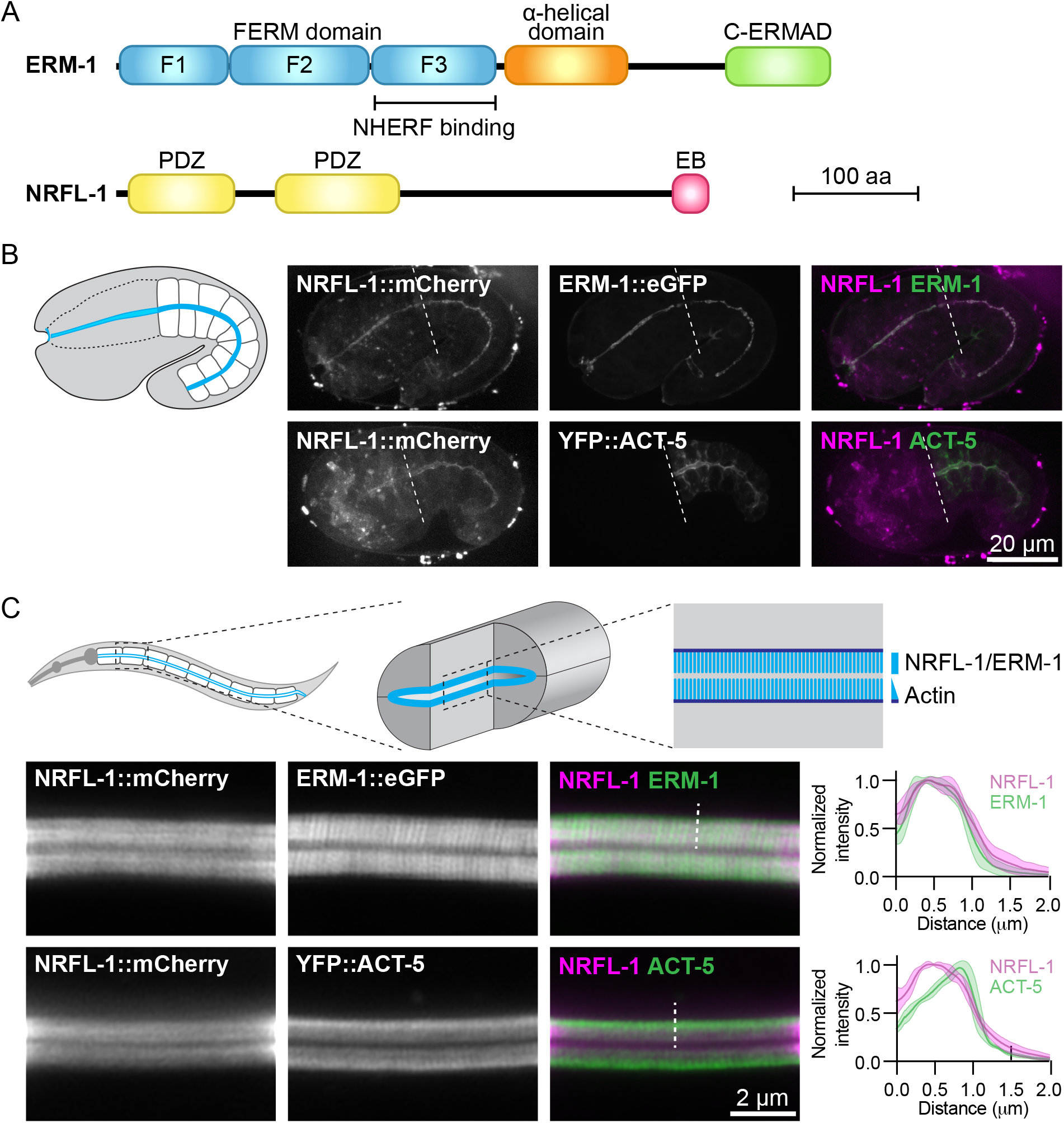
NRFL-1::mCherry localizes to the apical microvilli of intestinal cells. (**A**) Schematic representation of the domain organization of ERM-1 and NRFL-1. F1-F3 correspond to the three structural modules making up the FERM domain. FERM = Four-point-one, ezrin, radixin, moesin; C-ERMAD = C-terminal ezrin Radixin moesin (ERM) association domain; PDZ = Post-synaptic density-95, disks-large and zonula occludens-1; EB = ERM binding. (**B, C**) Distribution of NRFL-1::mCherry relative to ERM-1::GFP and YFP::ACT-5 in embryos (B) and apical membrane of L4 larval intestines (C). Dashed line in (B) separates the pharynx (left) from the intestine (right), and dashed line in (C) serves as an example of the line scan position used for the graphs on the right. Graphs plot the relative fluorescence intensity from the intestinal lumen to the cytoplasm. Solid line represents the mean and the shading lines the ± SD. n = 6 animals for both graphs. Images were taken using spinning-disk confocal (B) and Airyscan microscopes (C), and maximum intensity projections (B) or a single plane (C) are presented.

In contrast to ERM-1, little is known about the functioning of NRFL-1. A yeast-two hybrid screen identified the amino acid transporter family protein AAT-6 as an interactor of NRFL-1 (Hagiwara et al., 2012). However, the effects of NRFL-1 loss are minor. In aging adults, AAT-6 is no longer retained at the luminal membrane of the intestine in *nrfl-1* mutants, while younger *nrfl-1* mutants show increased mobility of AAT-6 by fluorescence recovery after photobleaching (FRAP). Moreover, *nrfl-1* mutants are homozygous viable, demonstrating that NRFL-1 is not critical for intestinal development (Hagiwara et al., 2012).

To investigate the relationship between ERM-1 and NRFL-1, we used CRISPR/Cas9 engineering to generate an *nrfl-1* deletion mutant, a mutant lacking the ERM-1 binding domain, and fluorescently tagged NRFL-1 variants. We show that NRFL-1 localizes to the apical microvillar domain of the intestine, and that this localization depends on the ability of NRFL-1 to bind to ERM-1 via the C-terminal ERM-1 binding domain. The loss of *nrfl-1* did not affect the localization, phosphorylation status, or protein dynamics of ERM-1, indicating that *C. elegans* NRFL-1 does not control the activity of ERM-1. However, when we combined the *nrfl-1* null mutant with *erm-1* mutants that block or mimic phosphorylation of the C-terminal threonine 544 residue, we observed severe intestinal defects, resembling the effects of strong loss of *erm-1* function. In mice, ezrin was shown to form distinct complexes with NHERF1 and actin. As the ERM-1 phosphorylation mutants affect the ability of ERM-1 to interact with actin, we postulate that the activities of ERM-1 in the intestine redundantly involve actin binding and the organization of apical domain composition through NRFL-1.

## Results

### NRFL-1 localizes to the apical domain through ERM-1 binding

To investigate the relationship between NRFL-1 and ERM-1, we first examined if NRFL-1 colocalizes with ERM-1. We used CRISPR/Cas9 to engineer an endogenous C-terminal NRFL-1::mCherry fusion, which tags all predicted isoforms. Animals homozygous for the *nrfl-1::mCherry* knock-in are viable and have a wild-type appearance. We detected expression of NRFL-1 in multiple epithelia including the intestine, excretory canal, pharynx, uterus, and spermatheca (Fig. 1B, C; Fig. S1B). In each of these tissues, NRFL-1::mCherry co-localized with an endogenous ERM-1::GFP fusion protein at the cortex (Fig. 1B, C; Fig. S1B). In the embryo, NRFL-1 localized to the nascent apical domain of intestinal cells, overlapping with ERM-1 and the epithelial actin ACT-5, visualized using a YFP::ACT-5 transgene (Fig. 1B). Confocal super resolution imaging of the intestine in larval stages showed co-localization of NRFL-1 with ERM-1::GFP and YFP::ACT-5 at microvilli, apical to the more intense belt of YFP::ACT-5 at the terminal web (Fig. 1C). The observed distribution of NRFL-1::mCherry is consistent with previous observations in *C. elegans* (Hagiwara et al., 2012), as well as with localization of EBP50 in mammalian epithelial tissues (Ingraffea, 2002; Morales et al., 2004; Kreimann et al., 2007).

We previously showed that ERM-1 and NRFL-1 interact in a yeast two-hybrid assay and in pull-downs from mammalian cultured cells (Koorman et al., 2016). To determine if these proteins interact in a more physiological setting, we used the recently developed split intein-mediated protein ligation (SIMPL) system that relies on protein splicing by split intein domains to detect protein–protein interactions (Yao et al., 2020). We ubiquitously expressed ERM-1 fused to the intein N-terminal fragment (IN) and the V5 epitope, and NRFL-1 fused to the C-terminal fragment (IC) and the FLAG epitope. We observed full splicing of NRFL-1 to ERM-1 by western blot of *C. elegans* lysates by observing a size shift and co-staining of the shifted band by both V5 and FLAG antibodies. In contrast, a negative control pair consisting of IC-tagged NRFL-1 and IN-tagged mKate2 showed only limited splicing of NRFL-1 to mKate2 (Fig. S2A). To visualize if splicing occurs *in vivo* in the intestine, we modified the SIMPL system by including a split mVenus tag. We added the mVenus N-terminal fragment (VN155) to ERM-1::V5-IN and mVenus C-terminal fragment (VC155) to IC-FLAG::NRFL-1, such that upon intein splicing the reconstituted mVenus becomes linked to NRFL-1 (Kodama and Hu, 2010). We readily observed localization of mVenus at the apical domain of intestinal cells, indicating that NRFL-1 and ERM-1 interact in this tissue (Fig. S2B).

We next investigated whether NRFL-1 distribution to the apical plasma membrane is dependent on ERM-1, by analyzing NRFL-1::mCherry upon tissue-specific depletion of ERM-1. To deplete ERM-1 in intestinal cells, we introduced an anti-GFP-nanobody::ZIF-1 fusion driven by the intestine-specific *elt-2* promoter as an extrachromosomal array in animals expressing endogenous ERM-1::GFP and NRFL-1::mCherry (Wang et al., 2017). Expression of the nanobody::ZIF-1 fusion resulted in variable levels of ERM-1::GFP depletion. The apical levels of ERM-1::GFP and NRFL-1::mCherry showed a linear correlation, indicating that apical recruitment of NRFL-1 in the intestine directly depends on ERM-1 (Fig. 2A).

**Figure 2.**
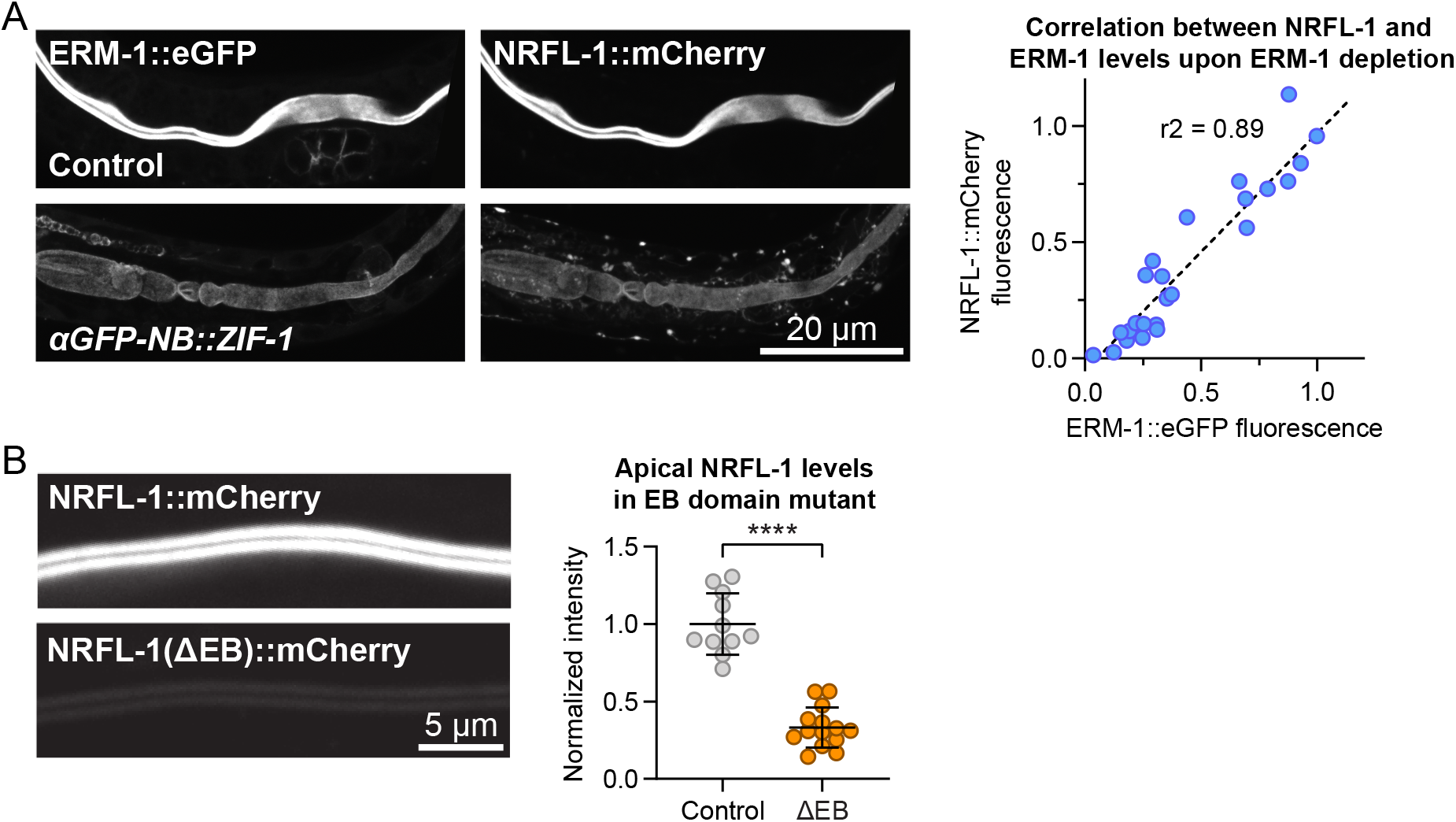
NRFL-1 localizes to the apical domain through ERM-1 binding. (**A**) Quantification of apical levels of NRFL-1::mCherry vs. ERM-1::GFP in L1 larval intestines upon different levels of ERM-1::GFP depletion by expression of an anti-GFP nanobody::ZIF-1 fusion protein. Fluorescence micrographs show representative examples, graph shows quantification of signal intensity at the apical membrane. Each data point in the graph represents a single animal, and the line a linear regression. Values are normalized to the mean intensity in control animals. n = 25 animals. (**B**) Quantification of apical levels of NRFL-1(ΔEB)::mCherry relative to NRFL-1::mCherry at the apical membrane of L1 larval intestines. Fluorescence micrographs show representative examples, and the graph the quantification. Each data point in the graph represents a single animal, and values are normalized to the mean intensity in control animals. Error bars: mean ± SD; Statistical test: Welch’s Student’s t-test; ^****^ = P ≤ 0.0001. n = 10 animals for NRFL-1::mCherry and 14 animals for NRFL-1(ΔEB)::mCherry. All images were taken using a spinning disk confocal microscope and presented as maximum intensity projections.

The interaction between mammalian EBP50 and ezrin requires the C-terminal EB domain (Reczek and Bretscher, 1998; Finnerty et al., 2004; Reczek et al., 1997), which is conserved in NRFL-1 (Fig. 1A; Fig. S1A). To determine if the NRFL-1 EB domain is required for the interaction with ERM-1, we repeated the SIMPL-mVenus experiment using an NRFL-1(ΔEB) mutant that lacks the C-terminal 28 amino acids of NRFL-1. Compared to wild-type NRFL-1, we observed only residual apical localization of mVenus in intestinal cells, indicating that the interaction of NRFL-1 with ERM-1 depends on the presence of the EB domain (Fig. S2B).

To determine if the EB domain is necessary for the apical localization of NRFL-1, we used CRISPR/Cas9 to engineer the 28 aa EB deletion in the *nrfl-1::mCherry* strain. The resulting *nrfl-1(Δeb)::mCherry* animals are homozygous viable, consistent with the lack of severe defects in previously described *nrfl-1* mutants (Hagiwara et al., 2012; Na et al., 2017). We detected a dramatic reduction in apical levels of NRFL-1(ΔEB)::mCherry in intestinal cells when compared with NRFL-1::mCherry (Fig. 2B). NRFL-1(ΔEB)::mCherry also failed to localize at the cortex in the uterus and spermatheca (Fig. S1C), while apical levels in the excretory canal were reduced (Fig. S1D). These results indicate that apical recruitment of NRFL-1 is mediated by the EB domain. However, the presence of some residual apical NRFL-1(ΔEB)::mCherry in the intestine and excretory canal suggests the existence of alternative membrane-targeting mechanisms. Collectively, our results show that the interaction between ERM and NHERF proteins is conserved in *C. elegans*, and that the localization of NRFL-1 is largely mediated by its interaction with ERM-1.

### NRFL-1 cooperates with ERM-1 phosphorylation in regulating intestinal lumen formation

We next wanted to investigate the effects of loss of NRFL-1 on intestinal lumen formation. Previous studies using partial deletion alleles of *nrfl-1* indicated that loss of NRFL-1 alone does not cause defects in the formation of the intestine (Hagiwara et al., 2012; Na et al., 2017). To rule out the possibility that the lack of severe defects is due to the production of truncated NRFL-1 proteins, we used CRISPR/Cas9 genome engineering to generate the *nrfl-1(mib59)* deletion allele. This allele lacks almost the entire *nrfl-1* locus and additionally causes a frameshift in the first exon of the long isoforms (Fig. 3A). Hence, we refer to *mib59* as *nrfl-1(null)*. Animals homozygous for the *nrfl-1(null)* allele are viable, have a healthy appearance, and normal brood sizes, confirming that NRFL-1 is not essential for *C. elegans* development (Fig. 3B).

**Figure 3.**
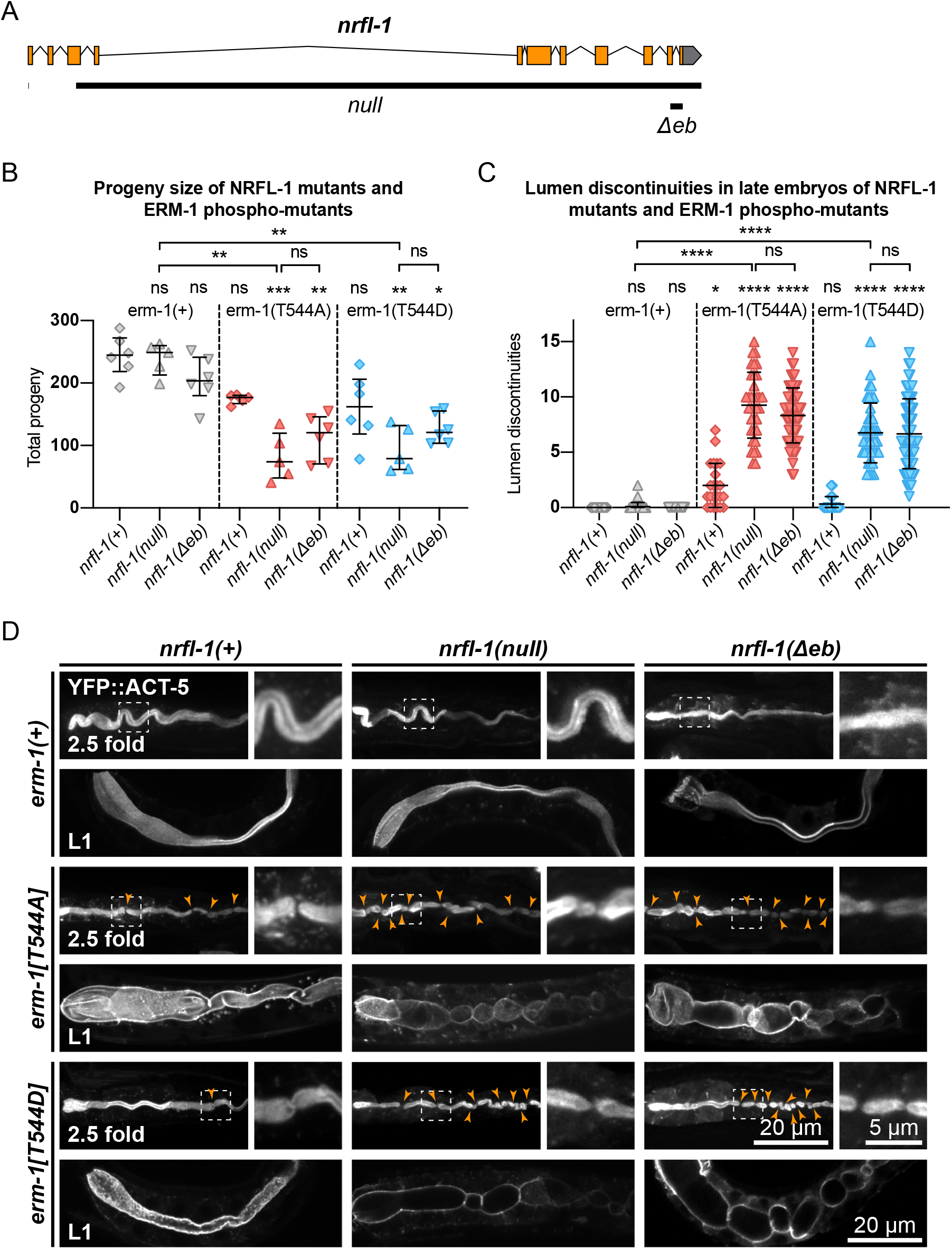
NRFL-1 cooperates with ERM-1 C-terminal phosphorylation in intestinal lumen formation. (A) Gene model for *nrfl-1a*. Orange boxes represent exons and lines represent introns. Grey box represents 3’ untranslated region. Black bars denote the regions deleted in *null* and *Δeb* alleles. (**B**) Quantification of total progeny from parents of indicated genotypes. Each data point represents the progeny of a single animal; N = 5 or 6. Error bars: mean ± SD. Statistical test: One-way Anova. (**C**) Quantification of lumen discontinuities in 2.5-fold stage embryos of indicated genotypes expressing YFP::ACT-5. Each data point represents a single animal. Error bars: mean ± SD. Statistical test: One-way Anova. *nrfl-1(+); erm-1(+)* n = 19, *nrfl-1(null); erm-1(+)* n = 35, *nrfl-1(Δebd); erm-1(+)* n = 48, *nrfl-1(+); erm-1[T544A]* n = 23, *nrfl-1(null); erm-1[T544A]* n = 35, *nrfl-1(Δebd); erm-1[T544A]* n = 61, *nrfl-1(+); erm-1[T544D]* n = 28, *nrfl-1(null); erm-1[T544D]* n = 41, *nrfl-1(Δebd); erm-1[T544D]* n = 69. (**D**) Representative images of intestinal defects in 2.5-fold stage embryos and L1 larvae of indicated genotypes, expressing YFP::ACT-5 as an apical marker. Images of the 2.5-fold stage embryos were computationally straightened, and the orange arrowheads indicate the constrictions in the lumen. All images are taken using a spinning-disk confocal microscope, and maximum intensity projections are presented.

One of the possible reasons for the lack of a severe intestinal phenotype in *nrfl-1(null)* animals is that NRFL-1 may only mediate part of the functions of ERM-1 in the intestine. To investigate this possibility, we made use of the non-phosphorylatable *erm-1[T544A]* and phosphomimetic *erm-1[T544D]* alleles we generated previously (Ramalho et al., 2020). Both mutants cause a delay in the apical recruitment of ERM-1 and actin during embryogenesis, and the appearance of constrictions along the course of the lumen that only occasionally persist to the L1 stage. In contrast to *erm-1* RNAi or strong loss-of-function alleles, however, these animals are viable. Thus, *erm-1[T544A]* and *erm-1[T544D]* represent partial loss-of-function alleles that may act as a sensitized background to reveal the contribution of NRFL-1 to ERM-1 functioning. Indeed, combining *nrfl-1(null)* with either *erm-1* phosphorylation mutant caused severe defects not observed in the single mutants. Double mutant animals had a developmental delay and strongly reduced brood size (Fig. 3B).

We next examined the formation of the intestinal lumen and actin distribution using YFP::ACT-5 as a marker. We did not detect any defects in apical enrichment of ACT-5 or intestinal morphology in *nrfl-1(null)* embryos and larvae (Fig. 3C, D). Combining the *erm-1[T544A]* and *erm-1[T544D]* alleles with the *nrfl-1(null)* allele significantly increased the frequency of intestinal constrictions and their persistence until larval development (Fig. 3C, D). Intestines of early larval *nrfl-1(null); erm-1[T544A]* and *nrfl-1(null); erm-1[T544D]* animals were characterized by a cystic appearance and multiple constrictions that block intestinal flow as seen in feeding assays with a fluorescent membrane-impermeable dextran (Fig. 3C, D; Fig. S3B). In surviving L2 or older animals, we only observed morphological defects but no lumen discontinuities, indicating that the early larval arrest in double mutants is due to a block of flow of food through the intestine (Fig 3D; Fig S3A).

Finally, as the EB domain is essential for the apical localization of NRFL-1 and its interaction with ERM-1, we determined if loss of the EB domain results in similar synergistic phenotypes with the ERM-1 phosphorylation mutants as complete loss of NRFL-1. We used CRISPR/Cas9 genome engineering to generate a second *nrfl-1(Δeb)* allele, also removing the final 28 aa but lacking the mCherry tag used above (Fig. 3A). Similar to our observations for the mCherry-tagged variant, homozygous *nrfl-1(Δeb)* mutants are viable and show no significant defects in brood size or intestinal development (Fig. 3B–D). However, when combined with *erm-1[T544A]* or *erm-1[T544D]*, the resulting double mutants showed similar defects in viability, growth, brood size, and intestinal development as observed using the *nrfl-1(null)* allele (Fig. 3B–D). Thus, the *nrfl-1(*Δ*eb)* allele behaves like a null allele of *nrfl-1*. Taken together, our data show that NRFL-1 and ERM-1 function together in promoting lumen formation in the *C. elegans* intestine, and the binding to ERM-1 is essential for the functioning of NRFL-1 in the intestine.

### NRFL-1 does not directly regulate ERM-1 activity

NRFL-1 could function together with ERM-1 in at least two ways. It could act as a scaffold protein that is required for ERM-1 to organize protein complexes at the membrane, or it could regulate the activity of ERM-1 itself. To distinguish between these possibilities, we investigated whether loss of NRFL-1 affects the distribution, mobility, or T544 phosphorylation status of ERM-1. We first analyzed the distribution of ERM-1::GFP in larval *nrfl-1(null)* mutants. We did not detect any change in ERM-1::GFP subcellular localization or levels at the apical membrane in the intestine (Fig. 4A). Moreover, FRAP analysis demonstrated that the mobility of ERM-1::GFP at the apical intestinal membrane was not significantly altered in *nrfl-1(null)* larvae (Fig. 4B). We next investigated whether NRFL-1 regulates ERM-1 C-terminal phosphorylation by staining *nrfl-1(null)* mutants with an antibody specific for the C-terminal phosphorylated form of ERM proteins (pERM). The residues used to raise this antibody are fully conserved between mammals and *C. elegans* (Ramalho et al., 2020). Nevertheless, we first confirmed the specificity of the antibody for T544 phosphorylated ERM-1 by immunostaining of ERM-1[T544A] mutant animals. We readily detected pERM staining of the intestinal lumen in wild-type larvae, while no staining was observed in ERM-1[T544A] animals (Fig. S4A). Moreover, treatment of embryos with a phosphatase abolished staining with the pERM antibody (Fig. S4B). Thus, the pERM antibody is specific for T544 phosphorylated ERM-1. We then stained *nrfl-1(+)* and *nrfl-1(null)* animals with the pERM antibody. In both backgrounds, the pERM antibody stained the lumen of the intestine, indicating that loss of *nrfl-1* does not significantly alter the phosphorylation status of the C-terminal regulatory threonine of ERM-1 (Fig. 4C). Taken together, our results show that NRFL-1 does not regulate the distribution, dynamics, or phosphorylation of ERM-1, and therefore does not seem to directly regulate ERM-1.

**Figure 4.**
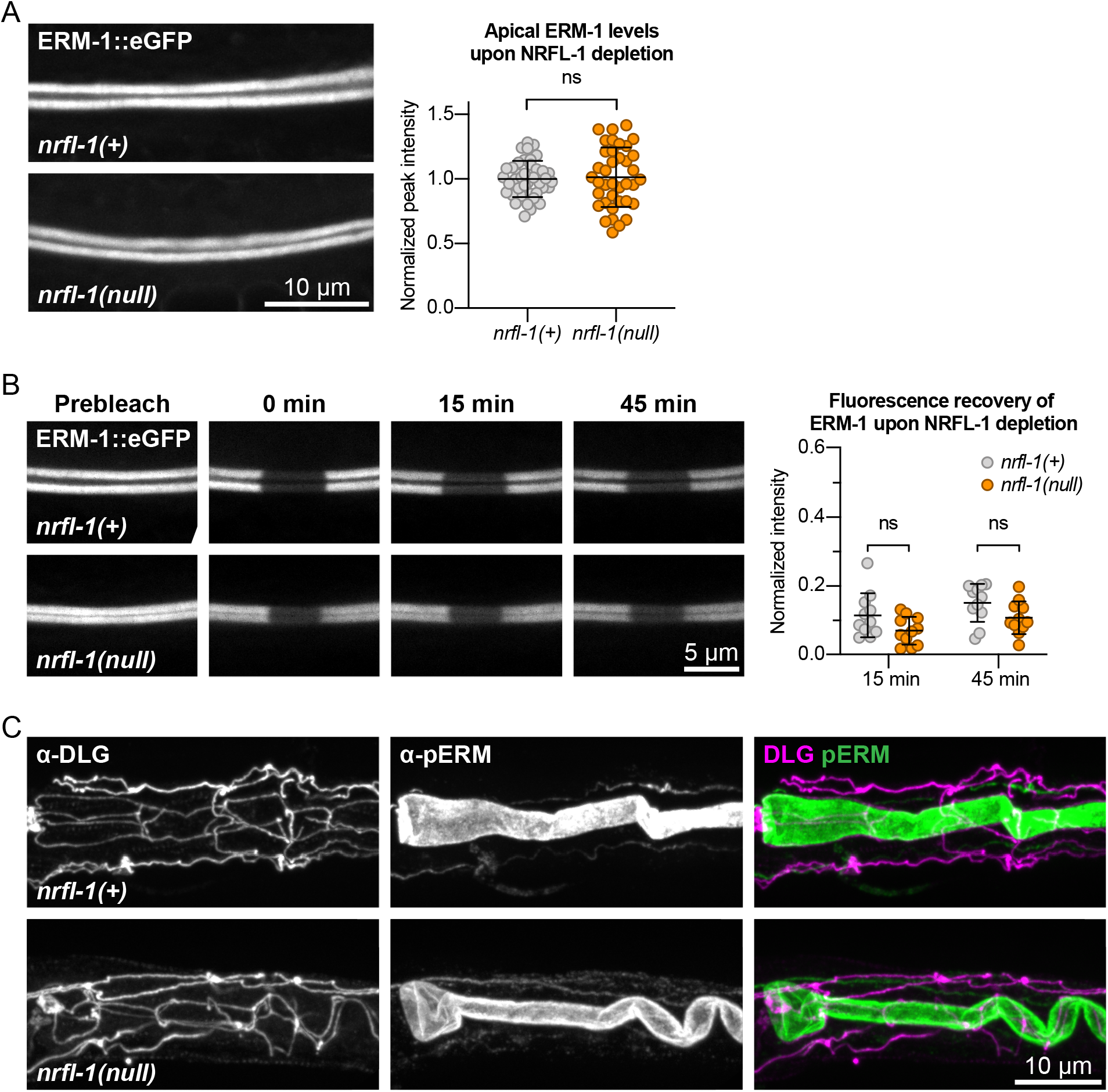
NRFL-1 does not regulate ERM-1 apical accumulation, dynamics, or phosphorylation status. (**A**) Representative images and quantification of ERM-1::GFP levels at the apical membrane of intestines in *nrfl-1(+)* and *nrfl-1(null)* L4 larvae. Each data point represents a single animal, and values are normalized to the mean intensity in control animals. Error bars: mean ± SD. Statistical test: Unpaired Student’s t-test. *nrfl-1(+)* n = 40, *nrfl-1(null)* n = 38. (**B**) FRAP analysis of apical ERM-1::GFP in the intestine of *nrfl-1(+)* and *nrfl-1(null)* L4 larvae. Fluorescence micrographs show representative examples. Graph shows the fluorescence intensity of ERM-1 in the photobleached region at the apical intestinal domain during recovery. Each data point represents a single animal, and values are relative to prebleach levels. Error bars: mean ± SD. Statistical test: Unpaired Student’s t-test. n = 11 for both genotypes and both timepoints. (**C**) Representative images of fixed *nrfl-1(+)* and *nrfl-1(null)* larvae stained with antibodies recognizing the junctional protein DLG-1 (α-DLG) and phosphorylated ERM-1 (α-pERM).

## Discussion

ERM and NHERF proteins function together in the specialization of polar membrane domains in several mammalian cell types. Here, we show that this cooperation is conserved in *C. elegans*, and that ERM-1 and NRFL-1 function together in lumen formation in the intestine. NRFL-1 physically interacts with ERM-1 through its C-terminal EB domain. The interaction with ERM-1 is responsible for the apical localization of NRFL-1 in the intestine, as depletion of ERM-1 or deletion of the EB domain result in an almost complete lack of apical NRFL-1::mCherry signal.

Somewhat surprisingly, loss of NRFL-1 by itself did not cause any noticeable defect in intestinal formation, animal development, or viability. Three previous partial deletion alleles of *nrfl-1* have been described: *ok2292, tm3501*, and *ok297* (Hagiwara et al., 2012, 1; Na et al., 2017). No severe defects in animal development were reported for *ok2292* or *tm3501* (Hagiwara et al., 2012). However, *nrfl-1(ok297)* animals were reported to have ruptured vulva and sterile phenotypes (Na et al., 2017). Given that neither of the other two previously characterized alleles nor our newly generated *nrfl-1* deletion allele display these phenotypes, we think it is likely that the *ok297* strain analyzed either contains additional background mutations or that *ok297* represents a neomorphic allele of *nrfl-1*.

The non-essential role of *nrfl-1* contrasts with data in mice, where NHERF1 loss causes defects in intestinal microvilli formation (Morales et al., 2004), and in *Drosophila*, where *Sip1* mutants cause morphological defects in the follicle cells surrounding the oocytes and late embryonic lethality (Hughes et al., 2010). However, combining an *nrfl-1(null)* deletion mutant with phosphorylation-defective *erm-1[T544A]* or *erm-1[T544D]* mutants resulted in severe defects in intestinal lumen formation. Double mutant animals often harbor multiple constrictions in the intestinal lumen and arrest during early larval development, likely due at least in part to the inability of luminal contents to travel through the digestive system.

The intestinal phenotype of *nrfl-1; erm-1[T544A/D]* double mutants is similar to the phenotype described for *erm-1(RNAi)* and the *erm-1(tm677)* deletion allele (Göbel et al., 2004; van Fürden et al., 2004). This suggests that the activity of ERM-1 in the intestine is mediated through phosphorylation of the T554 residue and recruitment of NRFL-1. As loss of T544 regulation and NRFL-1 binding act synergistically these two events likely represent separate activities of ERM-1. The exact consequences of altering T544 phosphorylation are not known, however, apical enrichment of the specialized actin ACT-5 is disrupted (Ramalho et al., 2020). This is in agreement with findings in other systems that C-terminal phosphorylation of ERM proteins is required for apical recruitment of actin (Abbattiscianni et al., 2016; Hipfner et al., 2004; Roch et al., 2010). Interestingly, fractionation experiments from kidney epithelial cells indicate that ERM proteins interact with actin and NHERF1 in distinct complexes (Morales et al., 2004). Together with the lack of ACT-5 defects in *nrfl-1* mutants, this presents a possible model for the activities of ERM-1 and NRFL-1 in which ERM-1 organizes the assembly of the apical actin cytoskeleton through binding to actin, and the recruitment or local distribution of membrane-associated proteins through the scaffolding activities of NRFL-1.

Finally, while NHERF proteins in other systems have been reported to affect the localization or activity of ERM-1 proteins, we found no evidence for such a role in *C. elegans*. The loss of NRFL-1 did not cause any noticeable defects in the localization or levels of ERM-1 at the apical membrane, in the mobility of ERM-1 as examined by FRAP, or in the phosphorylation of T544. Thus, NRFL-1 appears to function solely as an effector of ERM-1 in *C. elegans*. This difference may explain why loss of NHERF proteins can have similar phenotypes as loss of ERM in other systems, but not *C. elegans* (Broere et al., 2008; Garbett et al., 2010; Georgescu et al., 2014; Hughes et al., 2010; Morales et al., 2004). In organisms where NHERF loss affects the localization or activity of ERM proteins, the loss of NHERF would result in both a lack of protein scaffolding by NHERF and a reduction in actin organizing ability of ERM. Thus, the lack of a reciprocal relationship in *C. elegans* make it a unique model in which these different aspects of ERM protein function can be studied separately.

## Materials and methods

### *C. elegans* strains and culture conditions

*C. elegans* strains were cultured under standard conditions (Brenner, 1974). Only hermaphrodites were used, and all experiments were performed with animals grown at 15 °C or 20 °C on standard Nematode Growth Medium (NGM) agar plates seeded with OP50 *Escherichia coli*. Table 1 contains a list of all the strains used.

**Table 1.** List of *C. elegans* strains used.

| Strain | Genotype |
| --- | --- |
| N2 | Wild type |
| JM125 | *cals107[Pges-1::act-5::YFP]* |
| BOX163 | *erm-1(mib9[erm-1[p. T544D]] I* |
| BOX165 | *erm-1(mib10[erm-1[p. T544A]] I* |
| BOX196 | *erm-1(mib10[erm-1[p. T544A]] I;*<br>*cals107[Pges-1::act-5::YFP]* |
| BOX197 | *erm-1(mib9[erm-1[p.T544D]] I;*<br>*cals107[Pges-1::act-5::YFP]* |
| BOX273 | *mibls48[Pelt-2::TIR-1::tagBFP2-Lox511::tbb-2-3'UTR, IV:5014740-5014802 (cxTi10816 site))] IV* |
| BOX404 | *nrfl-1(mib59[nrfl-1a(null=c.-1_8del; c.287_1404+703))] IV* |
| BOX422 | *nrfl-1(mib73[nrfl-1::mCherry]] IV;*<br>*mibls48[Pelt-2::TIR-1::tagBFP2-Lox511::tbb-2-3'UTR, IV:5014740-5014802 (cxTi10816 site))] IV* |
| BOX428 | *erm-1(mib15[erm-1::eGFP]] I;*<br>*nrfl-1(mib73[nrfl-1::mCherry]] IV;*<br>*mibls48[Pelt-2::TIR-1::tagBFP2-Lox511::tbb-2-3'UTR, IV:5014740-5014802 (cxTi10816 site))] IV* |
| BOX429 | *nrfl-1(mib73[nrfl-1::mCherry]] IV;*<br>*mibls48[Pelt-2::TIR-1::tagBFP2-Lox511::tbb-2-3'UTR, IV:5014740-5014802 (cxTi10816 site))] IV;*<br>*cals107[Pges-1::act-5::YFP]* |
| BOX440 | *nrfl-1(mib75[nrfl-1a(Δeb=c.1318_1401del)::mCherry]] IV;*<br>*mibls48[Pelt-2::TIR-1::tagBFP2-Lox511::tbb-2-3'UTR, IV:5014740-5014802 (cxTi10816 site))] IV;*<br>*cals107[Pges-1::act-5::YFP]* |
| BOX597 | *nrfl-1(mib104[nrfl-1a(Δeb=c.1318_1401del))] IV* |
| BOX670 | *erm-1(mib10[erm-1[p.T544A]] I;*<br>*nrfl-1(mib59[nrfl-1a(null=c.-1_8del; c.287_1404+703))] IV* |
| BOX671 | *erm-1(mib9[erm-1[p.T544D]] I;*<br>*nrfl-1(mib59[nrfl-1a(null=c.-1_8del; c.287_1404+703))] IV* |
| BOX672 | *nrfl-1(mib59[nrfl-1a(null=c.-1_8del;*<br>*c.287_1404+703))] IV; cals107[Pges-1::act-5::YFP]* |
| BOX673 | *erm-1(mib10[erm-1[p.T544A]] I;*<br>*nrfl-1(mib59[nrfl-1a(null=c.-1_8del; c.287_1404+703))] IV;*<br>*cals107[Pges-1::act-5::YFP]* |
| BOX674 | *erm-1(mib9[erm-1[p.T544D]] I;*<br>*nrfl-1(mib59[nrfl-1a(null=c.-1_8del; c.287_1404+703))] IV; cals107[Pges-1::act-5::YFP]* |
| BOX675 | *erm-1(mib10[erm-1[p.T544A]] I; nrfl-1(mib104[nrfl-1a(Δeb=c.1318_1401del))] IV* |
| BOX676 | *erm-1(mib9[erm-1[p.T544D]] I; nrfl-1(mib104[nrfl-1a(Δeb=c.1318_1401del))] IV* |
| BOX677 | *nrfl-1(mib104[nrfl-1a(Δeb=c.1318_1401del))] IV; cals107[Pges-1::act-5::YFP]* |
| BOX678 | *erm-1(mib10[erm-1[p.T544A]] I;*<br>*nrfl-1(mib104[nrfl-1a(Δeb=c.1318_1401del))] IV;*<br>*cals107[Pges-1::act-5::YFP]* |
| BOX679 | *erm-1(mib9[erm-1[p.T544D]] I;*<br>*nrfl-1(mib104[nrfl-1a(Δeb=c.1318_1401del))] IV;*<br>*cals107[Pges-1::act-5::YFP]* |

**Table 2.**
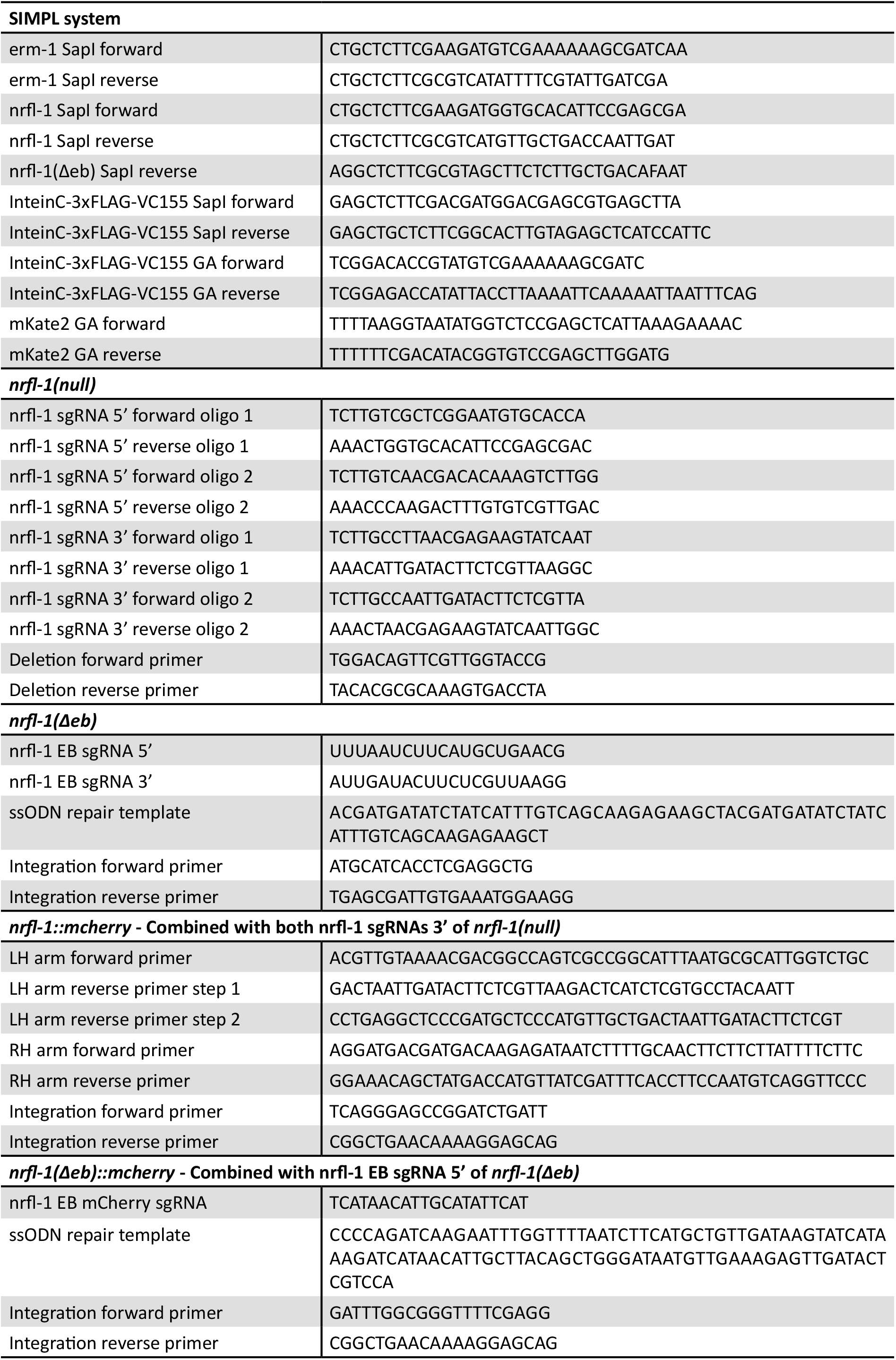
List of DNA and RNA sequences used.

### Cloning and strain generation for the SIMPL system

Bait and prey SIMPL constructs were generated using the SapI-based cloning strategy, as previously descripted (Yao et al., 2020). For the conventional SIMPL system, previously described intein inserts were used (Yao et al., 2020). For the SIMPL-mVenus system, the InteinC-3xFLAG-VC155 and VN155-HA-V5-InteinN inserts were codon-optimized for *C. elegans*, flanked by SapI sites and ordered as gBlocks (IDT). Primers containing the appropriate SapI overhangs were used to amplify *erm-1, nrfl-1* and *nrfl-1(Δeb)* from a cDNA library, InteinC-3xFLAG-VC155 from the ordered gBlock and mKate from pDD375 (Addgene #91825). All gBlocks and PCR products were blunt-end cloned into the plasmid pHSG298. Bait or prey, intein, the *rps-0* promoter and the *unc-54* 3’ UTR fragments were combined and inserted into the pMLS257 plasmid (Addgene #73716) using the SapTrap assembly method (Schwartz and Jorgensen, 2016; Yao et al., 2020). Finally for the SIMPL-mVenus system, an mKate2 sequence was integrated into the newly generated *Prps-0::ERM-1::VN155-HA-V5-InteinN::unc-54* plasmid. The mKate2 sequence and the *Prps-0::ERM-1::VN155-HA-V5-InteinN::unc-54* plasmid were amplified using primers with the appropriate overhangs to incorporate the mKate2 into the plasmid between the promotor and *erm-1* coding sequence using Gibson Assembly (GA). Constructs were verified by Sanger sequencing before injection (Macrogen Europe). Plasmids used for injection were purified using the PureLink HQ Mini Plasmid DNA Purification Kit (ThermoFisher) using the extra wash step and buffer recommended for endA+ strains. Final plasmid sequences are available in Genbank format in Supplemental File 1.

Transgenic animals expressing bait and prey constructs were generated by microinjection in the gonads of young adult N2 animals using an inverted microinjection setup (Eppendorf) with 20 ng/ μl of bait and prey plasmids, as well as the pDD382 plasmid (Addgene #91830) containing a visible dominant Rol marker and an hygromycin selection cassette. The DNA mix was spun at max speed on a tabletop centrifuge for 15 min prior to injection. Injected animals were incubated for 2–3 days at 20 °C before addition of hygromycin B (250 μg/ml) to the plates. After 1–2 days, surviving Rol animals were singled, allowed to develop, and F2 progeny was screened for successful transmission of the transgenic extrachromosomal array. Multiple lines with successful transmission were saved and used for analysis.

### Western blot SIMPL analysis

Animals were grown on NGM plates supplemented with hygromycin B (250 µg/ml) until plates were full, washed off with M9 buffer (0.22 M KH_2_PO, 0.42 M Na_2_HPO_4_, 0.85 M NaCl, 0.001 M MgSO_4_), washed three times with M9 buffer, and incubated at room temperature (RT) for 20 min. Samples were then pelleted and resuspended in 100–200 µl of lysis buffer (25 mM Tris-HCl pH 7.5, 150 mM NaCl, 1 mM EDTA, 0.5 % IGEPAL CA-630 (Sigma-Aldrich), 1 tablet/50 ml cOmplete protease inhibitor cocktail (Sigma-Aldrich)), and sonicated with a Diagenode BioRupter Plus for 10 min with the high setting and on/off cycles of 30 s in a 4 °C water bath. The lysates were spun at max speed for 15 min, an equal volume of 2× SDS buffer (100 mM Tris-HCl, 4 % SDS, 0.2% bromophenol blue, 20 % glycerol, and 10 % β -mercaptoethanol) was added, and boiled 10 min. Depending on the experiment, 5–12 µl of protein lysate was loaded into pre-cast protein gels (4–12 % Bolt Bis Tris Plus, ThermoFisher) together with 10 µl of the molecular marker (PageRuler prestained, ThermoFisher). Gels were run for 30–45 min at 200 V in NuPAGE MOPS SDS Running buffer (ThermoFisher), and transferred onto a PVDF membrane (Immobilon-P 0.45 µm, Millipore) at 4 °C and 30 V overnight in Bolt transfer buffer (ThermoFisher). For staining, membranes were rinsed in TBST (50 mM Tris-Cl, 150 mM NaCl, 0.1 % Tween-20), blocked with 4 % milk in TBST for 1 h at RT, and incubated with primary antibodies in milk for 1 h at RT. Membranes were washed three times for 10 min in TBST, incubated with secondary antibodies in milk for 1 h at RT, and washed again three times for 10 min in TBST before exposure using ECL (SignalFire Plus, Cell signaling). The following antibodies and concentrations were used: rabbit anti-V5, 1:1000 (Cell Signaling #13202); mouse anti-FLAG, 1:10000 (Sigma #F1804); goat anti-Rabbit and donkey anti-mouse HRP conjugates, 1:5000.

### CRISPR/Cas9 genome engineering

The *nrfl-1::mCherry, nrfl-1(Δeb)* and *nrfl-1(Δeb)::mCherry* strains were engineered by homology-directed repair of CRISPR/Cas9-induced DNA double-strand breaks (DSBs), while the *nrfl-1(null)* deletion was generated by imprecise repair of CRISPR/Cas9-induced DSBs. Delivery of components for CRISPR/ Cas9 editing was done by microinjection in the gonads of young adult animals of different genetic backgrounds: *nrfl-1(Δeb)* and *nrfl-1(mib59)* were generated in an N2 background; *nrfl-1::mCherry* was generated in BOX273 background, and *nrfl-1(Δeb)::mCherry* in a BOX422 background. All sequences of the oligonucleotides and crRNAs used (synthesized by IDT) are listed in Supplementary Table 2.

For the *nrfl-1::mCherry*, two plasmid-based sgRNAs were used, generated by ligation of annealed oligo pairs into the *pU6::sgRNA* expression vector pJJR50 (Addgene #75026) as previously described (Waaijers et al., 2016). To generate the *nrfl-1::mCherry* repair template we created a custom SEC vector, pJJR83 (Addgene #75028), by replacing a fragment of pDD282 (Addgene #66823) containing the GFP sequence with a similar fragment containing a codon optimized mCherry sequence with synthetic introns using the flanking Bsu36I and BglII restriction sites. Homology arms of about ±750 bp, flanking the DSB site, were amplified from genomic DNA and introduced into pJJR83 as previously described (Dickinson et al., 2015). The sgRNA (100 ng/μl) and SEC repair template (20 ng/μl) plasmids combined with *Peft-3::Cas9* (60 ng/μl; Addgene #46168) and *Pmyo-2::mCherry* co-injection marker (2.5 ng/μl; pCFJ90, Addgene #19327) were micro-injected in the gonad of young adults. Two injected animals were pooled per plate, incubated for 3 days at 20 °C, 500 μl of 5 mg/ml hygromycin was added per plate, and non-transgenic Rol animals were selected after 4–5 days. These selected animals were lysed and genotyped with primers flanking the homology arms and confirmed by Sanger sequencing. To eliminate the SEC selection cassette L1 progeny of homozygous Rol animals was heat shocked in a water-bath at 34 °C for 1 h.

To generate the *nrfl-1(null)* deletion allele a mix containing *Peft-3::Cas9* (Addgene #46168; 50 ng/μl), two pairs of sgRNA plasmids targeting the 5’ or 3’ ends of the *nrfl-1* open reading frame (75 ng/μl each), and a *dpy-10* sgRNA plasmid (50 ng/μl) for co-CRISPR selection (Arribere et al., 2014) were micro-injected in the gonad of young adults. To select for deletions, injected animals were transferred to individual plates, incubated for 3–4 days at 20 °C, and 96 non-transgenic F1 animals (wild-type, Dpy, or Rol) from 2–3 plates containing high numbers of Dpy and Rol animals were selected and transferred to individual plates. After laying eggs, F1 animals were lysed and genotyped with primers flanking the *nrfl-1* ORF. In all cases, deletions were confirmed by Sanger sequencing. Sanger sequencing was also used to determine the precise molecular lesion in selected animals. The *nrfl-1(null)* allele used in this paper, *nrfl-1(mib59)*, consists of a 9bp deletion starting 1 bp before the initial base of the start codon of long *nrfl-1* isoforms (a, c, d, h, j), and a second 11,537 bp deletion spanning part of the third exon (791 bp from start of *nrfl-1a*) until the downstream intergenic region, which includes the entire ORFs of the small *nrfl-1* isoforms (left flank 5’-atgcttgtgatctctgaagaaggag, right flank 5’ aatatcacgaacaacttctaggagc). The *mib59* allele also deleted an ncRNA (C01F6.16) and three piRNAs (C01F6.10, F32B2.25, and F23B2.28) located within *nrfl-1* introns.

The NRFL-1 EB domain deletions were generated using the Alt-R CRISPR-Cas9 system (IDT). A single-stranded oligodeoxynucleotide with about 35 bp homology arms was used as a repair template to fuse the flanks of a deletion spanning nucleotides 1390–1473 of *nrfl-1h*, as previously descripted (Dokshin et al., 2018). A mix of 250 ng/μl Cas9 protein, 2 μM repair template, 4.5 μM each *nrfl-1* crRNAs, 10 μM tracrRNA, as well as 1 μM *dpy-10* crRNA and ssODN repair for co-CRISPR selection (Arribere et al., 2014) was micro-injected into the gonads of young adults. Animals were selected as described above for the *nrfl-1(null)* allele and genotyped using two primers flanking the deletion.

### Microscopy and image analysis

Imaging of *C. elegans* was done by mounting embryos or larvae on a 5 % agarose pad in 20 mM Tetramisole solution in M9 to induce paralysis. Spinning disk confocal imaging was performed using a Nikon Ti-U microscope equipped with a Yokogawa CSU-X1 spinning disk using a 60×-1.4 NA objective, 488 nm and 561 nm lasers, Semrock “530” GFP-L and “600” TxRed emission filters, and an Andor iXON DU-885 camera. Imaging for FRAP and immunohistochemistry experiments was performed on a Nikon Eclipse-Ti microscope equipped with a Yokogawa CSU-X1-A1 spinning disk using 60× and 100× 1.4 NA objectives, ET-DAPI (49000), ET-GFP (49002), ET-mCherry (49008) emission filters, 355 nm, 488 nm, 491 nm, and 561 nm lasers, and a Photometrics Evolve 512 EMCCD camera. Targeted photobleaching was done using an ILas system (Roper Scientific France/ PICT-IBiSA, Institut Curie). Spinning disk images were acquired using MetaMorph Microscopy Automation & Image Analysis Software. All stacks along the z-axis were obtained at 0.25 μm intervals. Super resolution images of the microvilli were obtained using a Zeiss AxioObserver 7 SP microscope operated by Zeiss ZEN software with an Airyscan module GaAsP using a 100× 1.46 NA objective, 488 nm and 561 nm lasers, and Spectral 3 PMT detector. Maximum intensity Z projections were done in ImageJ (Fiji) software (Rueden et al., 2017; Schindelin et al., 2012). For quantifications, the same laser power and exposure times were used within experiments. Image scales were calibrated for each microscope using a micrometer slide. For display in figures, level adjustments, false coloring, and image overlays were done in Adobe Photoshop. Image rotation, cropping, and panel assembly were done in Adobe Illustrator. All edits were done non-destructively using adjustment layers and clipping masks, and images were kept in their original capture bit depth until final export from Illustrator for publication.

### Quantitative image analysis

Quantitative analysis of spinning disk images was done in Fiji. All values were corrected for background levels by subtracting the average of three regions within the field-of-view that did not contain any animals. For the apical levels, measurements were done in intestinal cells forming int2 through int6, and where the opposing apical membranes could be clearly seen as two lines. Levels were obtained by averaging the peak values of intensity profiles from three 25 px-wide (10 px-wide for the SIMPL-mVenus system) line-scans perpendicular to the membrane per animal. Intensity distribution profiles to analyze co-distribution of NRFL-1 with ERM-1 and ACT-5 were obtained by taking three 25 px wide line-scans perpendicular to the apical membrane in each animal. Before averaging these three values, they were aligned and normalized to the peak value. Measurements of multiple animals were again aligned based on the peak value. All presented graphs were made using GraphPad Prism and Adobe Illustrator.

### Protein degradation

For protein degradation using the anti-GFP-nanobody::ZIF-1 approach (Wang et al., 2017), gonads of young adult BOX428 animals were microinjected with 30 ng/μl *Pelt-2::*α*GFP-NB::ZIF-1* and 2.5 ng/μl *Pmyo-2::GFP* (#Addgene 26347) as a co-injection marker. Transgenic F1 animals were transferred to individual plates, F2 progeny was screened for successful transmission of the extrachromosomal array and imaged using spinning disk microscopy.

### Brood size

L4 animals were put on individual plates at 20 °C and transferred to a new plate daily until they died. After the parent was removed from a plate, hatched animals and the unhatched eggs were counted 2–4 days later. The number of animals and unhatched eggs combined constitutes the total progeny size. The graph presented was made using GraphPad Prism and Adobe Illustrator.

### Texas red-dextran assay

Mixed stage populations were collected in M9 and washed two times in M9. Animals were then pelleted, concentrated, resuspended in 1 mg/ml Texas Red-dextran 40,000 MW (Thermofisher D1829) in egg buffer (118 mM NaCl, 48 mM KCl, 2 mM MgCl2, 2 mM CaCl2, 25 mM HEPES pH 7.3), and incubated for 60 min on a shaker at 500 rpm. The dye in solution was removed by washing the samples with M9 two times. Animals were paralyzed in 10 mM Tetramisole, transferred to an agarose pad on a glass slide, and imaged using spinning disk microscopy.

### FRAP experiments and analysis

For FRAP assays, laser power was adjusted in each experiment to avoid complete photobleaching of the selected area, as the time scale of experiments prevented assessment of photo-induced damage. Photobleaching was performed on a circular region with a diameter of 30 or 40 px at the cortex, and images were taken just before bleaching, directly after, after 15 min, and after 45 min. These images were analyzed using ImageJ. The size of the area for FRAP analysis was defined by the full width at half maximum of an intensity plot across the bleached region. For each time point, the mean intensity value within the bleached region was determined, and the background, defined as the mean intensity of a non-bleached region outside the animal, was subtracted. The mean intensities within the bleached region were corrected for acquisition photobleaching per frame using the background-subtracted mean intensity of a similar non-bleached region at the cortex, which was normalized to the corresponding pre-bleach mean intensity. FRAP recovery was calculated as the change in corrected intensity values within the bleach region form the first image after bleach normalized to the mean intensity of the just before bleaching.

### Immunohistochemistry

For the staining of larval stages, embryos were obtained from gravid adults by bleaching and allowed to hatch and develop on plates at 15 °C for 24 h. Animals were collected from plates and washed three times with M9 and once with MQ H_2_O before being transferred to poly-L-lysine-coated frosted slides. For the staining of embryos, embryos were obtained from gravid adults by dissection in MQ H_2_O on poly-L-lysine-coated frosted slides and allowed to develop at RT for 4 h. A coverslip (Carl Roth, #1) was lowered on top of larvae/embryos, followed by freezing in liquid nitrogen and snapping off of the coverslip. Fixation was performed in formaldehyde solution with phosphatase inhibitors (3,7% formaldehyde (Sigma-Aldrich), 250 µM EDTA and 50 mM NaF in PBS (1,35 M NaCl, 27 mM KCl, 100 mM Na_2_HPO_4_, 18 mM KH_2_PO_4_)) at RT for 10 min. Samples were rinsed in PBS, permeabilized (PBS + 0,5% triton X-100 (Sigma-Aldrich)) for 30 min, washed four times in wash buffer (0,1% Triton X-100, 250 µM EDTA and 50 mM NaF in PBS) for 10 min each and then blocked (1% bovine serum albumin (Sigma-Aldrich) and 10% goat serum (Sigma-Aldrich)) for 1 h at RT. For the staining with protein phosphatase treatment, samples were treated with Lambda Protein Phosphatase (NEB) for 30 min at 30 °C followed with an additional four times washing step before they were blocked. Primary antibodies (anti-phospho-ezrin (Thr567)/radixin (Thr564)/moesin (Thr558) (48G2) rabbit mAb #3726 (Cell Signaling Technologies) 1:200 and mouse anti-DLG (Hybridoma bank) 1:50) in blocking solution were applied overnight at 4 °C. Samples were then washed four times in wash buffer for 10 min each and stained with secondary antibodies (Alexa-Fluor 488 goat anti-rabbit and Alexa-Fluor 568 goat anti-mouse (Life Technologies, A-11008 and A11004), both 1:500) in blocking solution for one hour at RT. Samples were then washed four times in wash buffer and once in PBS for 10 min each and finally mounted with Prolong Gold Antifade with DAPI (Thermofisher) under a coverslip and sealed with nail polish.

### Statistical analysis

All statistical analyses were performed using GraphPad Prism 8. For population comparisons, a D’Agostino & Pearson test of normality was first performed to determine if the data was sampled from a Gaussian distribution. For data drawn from a Gaussian distribution, comparisons between two populations were done using an unpaired t test, with Welch’s correction if the SDs of the populations differed significantly, and comparisons between >2 populations were done using a one-way ANOVA, or a Welch’s ANOVA if the SDs of the populations differed significantly. For data not drawn from a Gaussian distribution, a non-parametric test was used (Mann-Whitney for 2 populations and Kruskal-Wallis for >2 populations). ANOVA and non-parametric tests were followed up with multiple comparison tests of significance (Dunnett’s, Tukey’s, Dunnett’s T3 or Dunn’s). Tests of significance used and sample sizes are indicated in the figure legends. No statistical method was used to pre-determine sample sizes. No samples or animals were excluded from analysis. The experiments were not randomized, and the investigators were not blinded to allocation during experiments and outcome assessment.

## Acknowledgements and funding

We thank V. Portegijs and S. van den Heuvel for the *Pelt-2::*α*GFP-NB::ZIF-1* plasmid, H.R. Pires for strain BOX273, and members of the S. van den Heuvel, S. Ruijtenberg, and M. Boxem groups for helpful discussions. We also thank Wormbase (Harris et al., 2020) and the Biology Imaging Center, Faculty of Sciences, Department of Biology, Utrecht University. Some strains were provided by the Caenorhabditis Genetics Center, which is funded by NIH Office of Research Infrastructure Programs (P40 OD010440). This work was supported by the Netherlands Organization for Scientific Research (NWO)-ALW Open Program 824.14.021 and NWO-VICI 016.VICI.170.165 grants to M. Boxem.

## Author contributions

Conceptualization: J.J.S., J.J.R., M.B.; Methodology: J.J.S., J.J.R., J.R.K., R.S.; Formal analysis: J.J.S., J.J.R., J.R.K.; Investigation: J.J.S., J.J.R., J.R.K., R.S.; Writing - original draft: J.J.S., J.J.R., M.B.; Writing – review & editing: J.R.K., R.S.; Visualization: J.J.S., J.J.R.; Supervision: M.B.; Project administration: M.B.; Funding acquisition: M.B.

---

**Figure S1.**
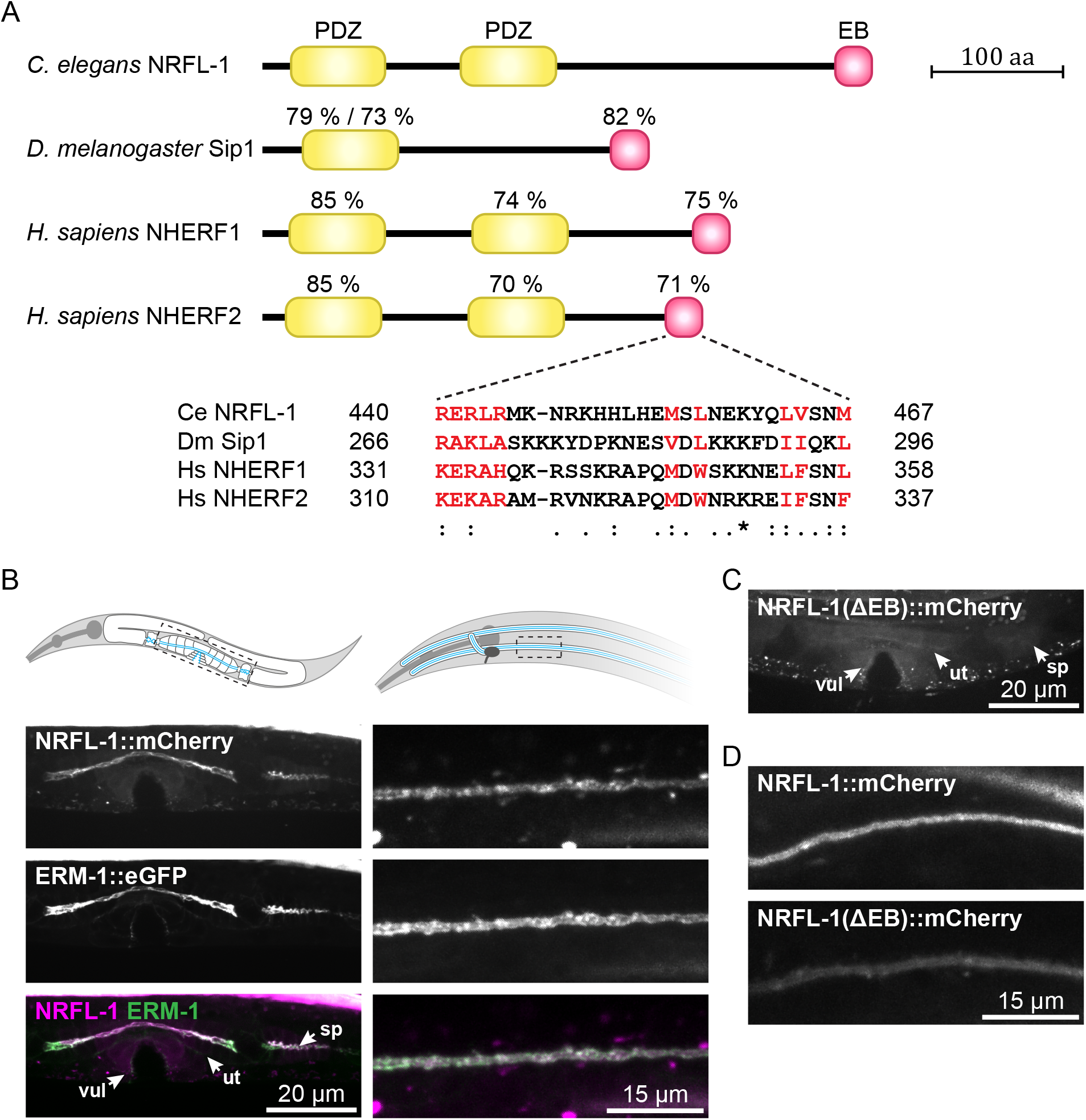
NRFL-1 is recruited to the apical domain by ERM-1 in different tissues. (**A**) Schematic representation of the domain organization of *C. elegans* NRFL-1, *D. melanogaster* Sip1 and *H. sapiens* NHERF1 and NHERF2. Percentages above the domains represent the similarity between that domain and the corresponding domain of NRFL-1. For the single PDZ domain of Sip1, two percentages are presented corresponding to each NRFL-1 PDZ domain. In the EB domain alignment, amino acids that are important for the interaction with ERM proteins are shown in red (Terawaki et al., 2006). PDZ = Post-synaptic density-95, disks-large and zonula occludens-1; EB = ERM binding. (**B**) Distribution of NRFL-1::mCherry and ERM-1::GFP in the excretory canal in L1 larvae (left panels), and in the vulva (vul), uterus (ut) and spermatheca (sp) in L4 larvae (right panels). (**C**) Distribution of NRFL-1(ΔEB)::mCherry in the vulva, uterus, and spermatheca of L4 larvae (compare with left panels in B). (**D**) Localization of NRFL-1::mCherry and NRFL-1(ΔEB)::mCherry in the excretory canal. Images of the same tissue were acquired and displayed with the same settings for comparison. All images are taken using a spinning-disk confocal microscope and maximum intensity projections are presented.

**Figure S2.**
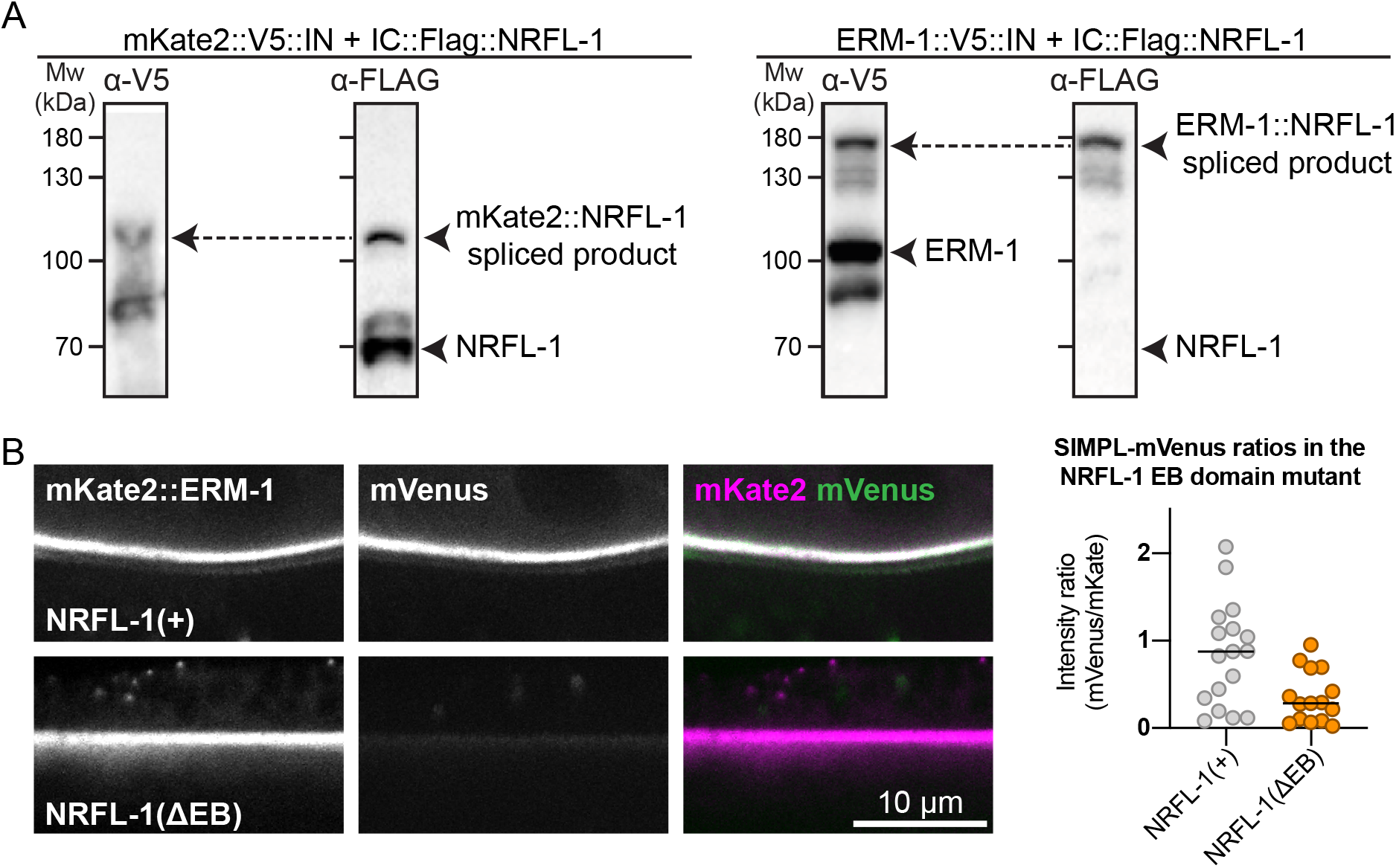
NRFL-1 interacts with ERM-1 *in vivo*. (**A**) Detection of an ERM-1–NRFL-1 interaction using the SIMPL system. V5 and FLAG epitopes are detected by western blot. Arrowheads indicate both unspliced proteins and the higher molecular weight covalently linked fusion proteins, generated by Intein splicing activity. Little splicing of NRFL-1 is observed with the control mKate2::V5::IN protein, while all NRFL-1 is spliced to ERM-1 in animals expressing ERM-1::V5::IN (**B**) Detection of an interaction of ERM-1 with wild-type NRFL-1, but not with NRFL-1(ΔEB), using the SIMPL-mVenus system. NRFL-1a::InteinC-3xFLAG-VC155 [NRFL-1(+)] or NRFL-1a(ΔEB)::InteinC-3xFLAG-VC155 [NRFL-1(ΔEB)] are expressed with mKate2::ERM-1::VN155-HA-V5-InteinN (mKate2::ERM-1). Fluorescence micrographs show representative examples. Graphs show quantification of apical mVenus levels, expressed as a ratio over mKate2::ERM-1 to account for varying expression levels of the extrachromosomal array. Each data point represents a single intestinal cell. Lines indicate median. N = 17 cells for NRFL-1(+) and 15 cells for NRFL-1(ΔEB). All images were taken using a spinning-disk confocal microscope and a single plane is presented.

**Figure S3.**
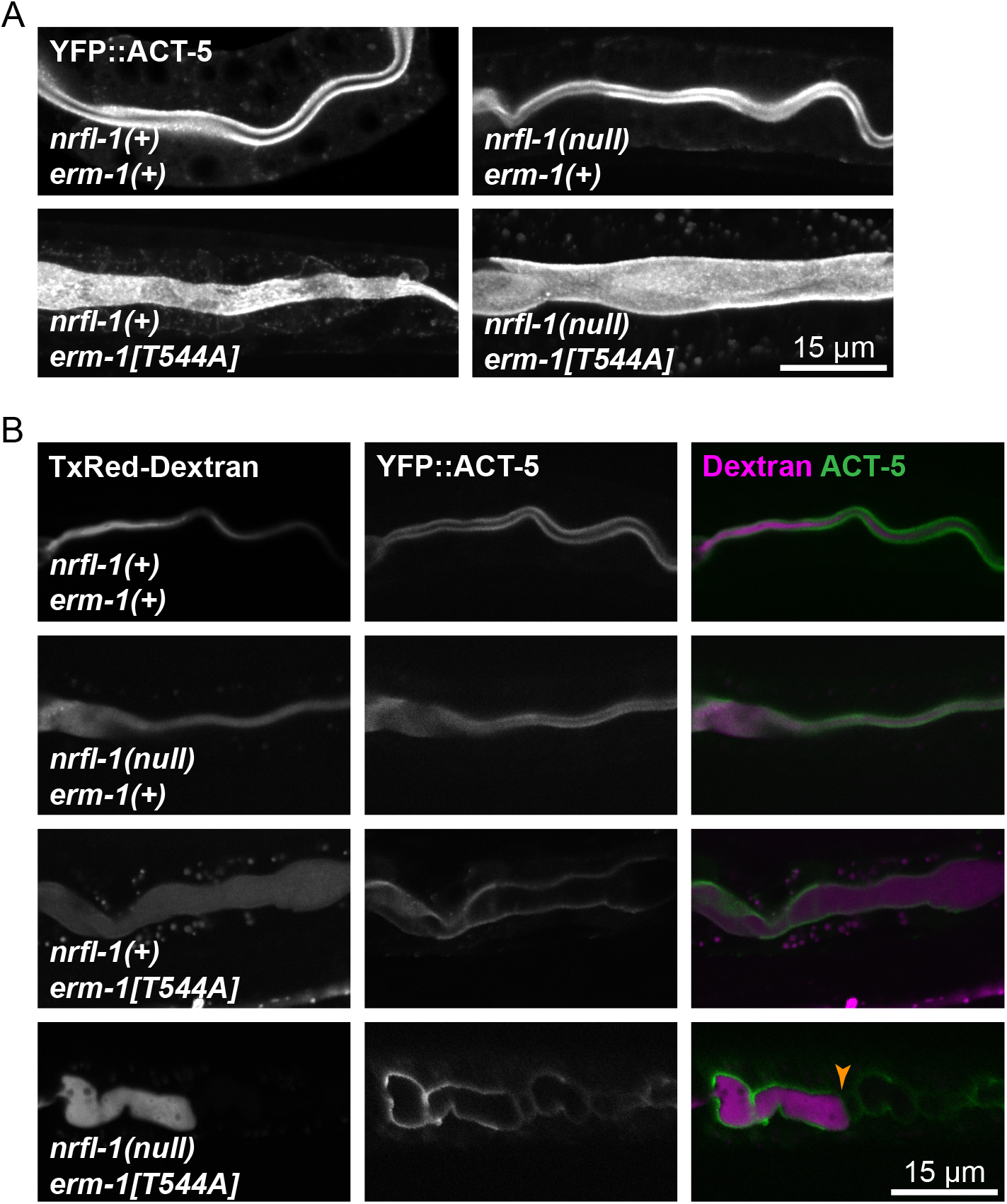
Lumen discontinuities in *nrfl-1* and *erm-1* single and double mutants. (**A**) Representative images of the intestine in L2 larvae of the indicated genotypes expressing YFP::ACT-5 as an apical marker. (**B**) L1 larvae carrying the apical marker YFP::ACT-5 of the indicated genotypes fed with Texas-Red Dextran. The orange arrowhead indicates the constriction that prevents the flow of fluorescent dye along the intestine. All images are taken using a spinning-disk confocal microscope, and a single focal plane is shown.

**Figure S4.**
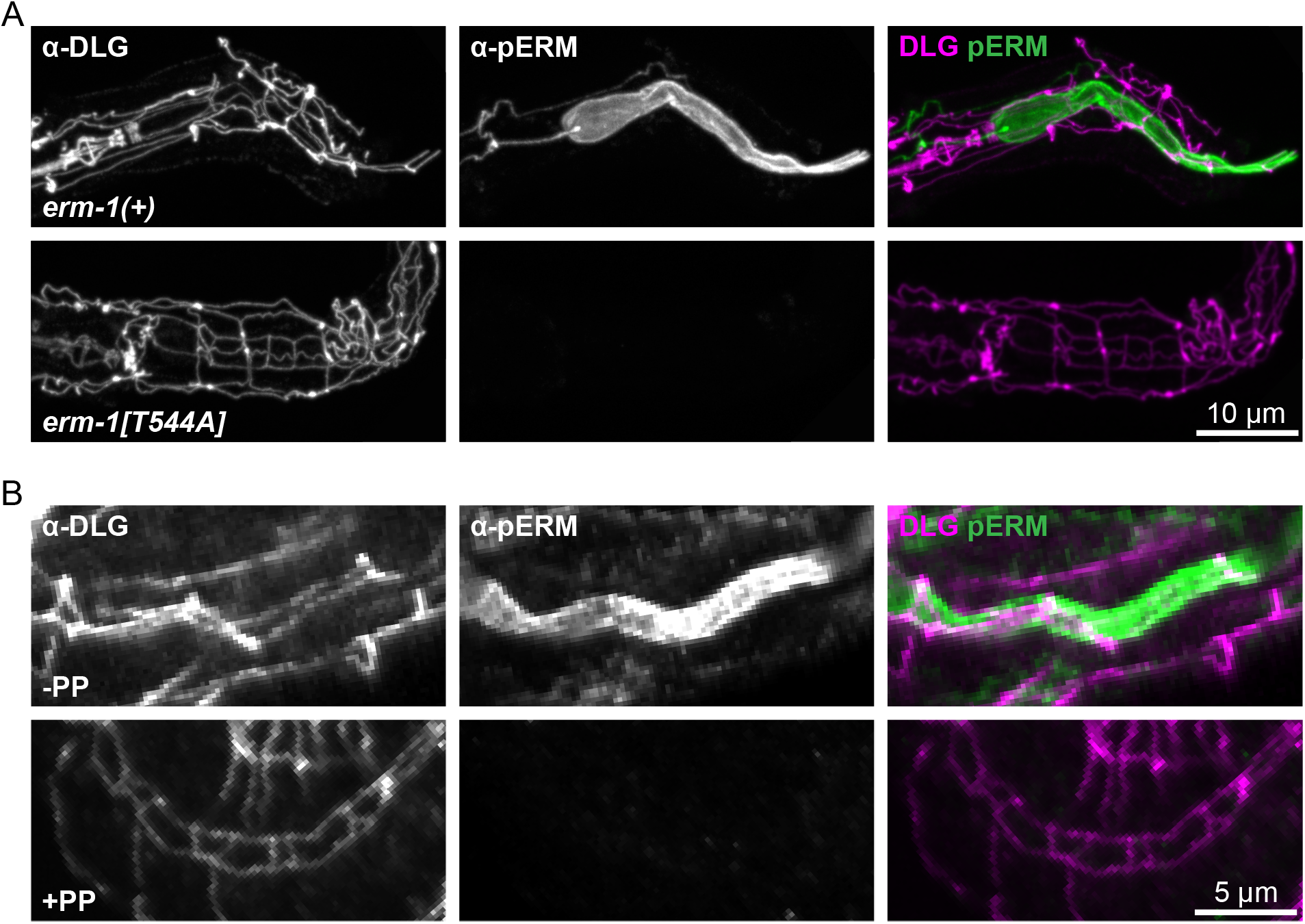
Validation of the specificity of the α-pERM for T544 phosphorylated ERM-1. (**A, B**) Representative images of fixed animals stained with antibodies recognizing the junctional protein DLG-1 (α-DLG) and phosphorylated ERM-1 (α-pERM). (A) shows *erm-1(+)* and *erm-1[T544A]* larvae and (B) shows the intestine of wild-type 2.5-fold embryos that are untreated (-PP) and treated with protein phosphatase (+PP). All images are taken using a spinning-disk confocal microscope, and maximum intensity projections are presented.

## Notes

### Competing Interest Statement

The authors have declared no competing interest.

